# On the correspondence of electrical and optical physiology in *in vivo* population-scale two-photon calcium imaging

**DOI:** 10.1101/800102

**Authors:** Peter Ledochowitsch, Lawrence Huang, Ulf Knoblich, Michael Oliver, Jerome Lecoq, Clay Reid, Lu Li, Hongkui Zeng, Christof Koch, Jack Waters, Saskia E.J. de Vries, Michael A. Buice

## Abstract

Multiphoton calcium imaging is commonly used to monitor the spiking of large populations of neurons. Recovering action potentials from fluorescence necessitates calibration experiments, often with simultaneous imaging and cell-attached recording. Here we performed calibration for imaging conditions matching those of the Allen Brain Observatory. We developed a novel crowd-sourced, algorithmic approach to quality control. Our final data set was 50 recordings from 35 neurons in 3 mouse lines. Our calibration indicated that 3 or more spikes were required to produce consistent changes in fluorescence. Moreover, neither a simple linear model nor a more complex biophysical model accurately predicted fluorescence for small numbers of spikes (1-3). We observed increases in fluorescence corresponding to prolonged depolarizations, particularly in Emx1-IRES-Cre mouse line crosses. Our results indicate that deriving spike times from fluorescence measurements may be an intractable problem in some mouse lines.

## Introduction

Systems neuroscience demands a steady increase in the number of simultaneously recorded neurons. Over the last five decades, the number of recorded neurons has doubled approximately every 7 years, mimicking Moore’s Law [1]. Electrophysiology has been the *de facto* gold standard for measuring neural activity for more than 80 years, starting with pioneering work on the squid giant axon by Young, Curtis, Cole, Hodgkin, and Huxley [2–4]. However, when the express goal is to push the envelope on the number of simultaneously recorded neurons *in vivo*, intracellular electrophysiology is impractical, and extracellular electrophysiology has other limitations, e.g. that the precise origin of the signal (positional, genetic, morphological, and physiological details of the generating neuron) are typically unknown [5]. Simultaneous recordings from large numbers of genetically-identified neurons or experiments that necessitate recording from the same neurons repeatedly over many days are difficult to achieve with electrophysiology and are more tractable with imaging techniques.

One commonly used imaging technique is 2-photon calcium imaging. Anticipated from first principles by Maria Göppert-Mayer [6, 7], and experimentally pioneered by Denk, Strickler, and Webb [8], 2-photon imaging emerged as a complementary technique for the acquisition of neural activity with single-cell resolution from hundreds of neurons in the living brain. Modern technology has enabled the development of genetically encoded calcium indicators (GECIs), which can be introduced via viral transfection or transgenic strategy, and are expressed in a promoter-specific manner. The development of GCaMP6 in particular, with its promise of elusive single spike sensitivity [9], heralded a new era of massive optophysiological surveys [10], recording large numbers of neurons from specific, genetically distinct populations.

Unfortunately, the fluorescence of a calcium indicator is only indirectly linked to spiking activity. For example, neurons typically exhibit large inward calcium currents during the falling phase of the action potential [11], but the details can be quite complex. A plethora of intracellular processes other than action potentials rely on calcium signaling and there is no guarantee that, even in the same neuron, each spike will be associated with the same distribution of calcium currents. Moreover, binding interactions between calcium, intracellular calcium buffers, and the indicator are complex, with cooperative binding at multiple sites on GCaMP indicators [12], leading to fluorescence that is dependent on the instantaneous intracellular calcium concentration and also on the firing history of the neuron.

The time resolution of two photon imaging is subject to technological limitations as well. The need to collect enough photons results in a trade-off between spatial and temporal resolution; practical sampling rates of ≈ 30Hz tend to vastly undersample the characteristic time scales of action potentials — 0.5-2 ms). The time course of calcium activity is further subject to fundamental biochemical constraints (e.g. calcium-indicator binding kinetics). This non-exhaustive list of caveats and complications highlights the need for accurate calibration experiments that quantify the relationship between action potentials and calcium activity.

Calibration against the gold standard of electrophysiology has often been performed during the initial characterization of indicators’ properties, e.g. Chen et al [9]. These calibration data are usually reported from virally infected neurons and using a small (single-cell-sized) field of view to maximize the signal-to-noise ratio. Huang et al [13] recently presented comparable calibration results from several GCaMP6 transgenic mouse lines, mostly imaged with a small field of view (20*μm* × 20*μm*, just large enough to contain a single neuron). This **in vivo** data is from 91 visually stimulated transgenic mouse neurons (L2/3, primary visual cortex) across several Cre-defined cell types (expressing GCaMp6f under Emx1 and Cux2 promoters, and GCaMp6s under Emx1 and tetO promoters) and includes simultaneously acquired, cell-attached electrophysiology. While Chen et al. showed single-spike detection to be feasible under their chosen expression and imaging conditions, Huang et al reported comparable fluorescence transients for GCaMP6f but smaller responses for GCaMP6s transgenics. In Chen et al, the 2P data were acquired at spatiotemporal resolutions much higher than what is typically achieved in population-level optophysiological studies (except for recent advances, which require their own calibration, e.g. [14]). Here we extend the results of Huang et al [13] to calibrate imaging conditions comparable to many population imaging studies by resampling the recordings both spatially and temporally to match the noise profile of the Allen Brain Observatory.

In addition to the problem of calibration, neuroscience, like many fields, faces an issue of lacking standardization and repeatability, which can be ameliorated by systematic, semi-automated quality control of research data, which helps to avoid the human bias typically encountered. Quality control of the data in [13] was conducted, as is common in the field, manually, compared to a more automated quality-control stage in large surveys like the Allen Brain Observatory [10]. As part of this work, we outline an approach for selecting recordings for analysis by constructing an algorithm that curates for recording features characteristic of traces selected by multiple, independent expert annotators. This approach aims to achieve greater transparency and reproducibility by removing a significant degree of subjectivity from analysis.

Based on the obtained calibration data set, we assessed, through a combination of analysis and modeling, the likelihood that a given number of action potentials would leave an algorithmically detectable ‘event’ in the fluorescence trace. First we asked whether we can quantify the correspondence between identified calcium events (as would be derived from a “spike” inference algorithm) and spiking activity. Second, we investigated to what extent models of calcium can generate observed fluorescence from spike trains. We show that in conditions of low instantaneous firing rates, spikes can be difficult or impossible to infer from fluorescence and, conversely, fluorescence can be difficult to predict from spiking activity. We argue that in some neurons, the variance in fluorescence is more readily explained by electrophysiological features other than the action potentials the neuron generated, and that these features are Cre-line dependent.

## Materials and Methods

### The data set

We start with data from the same set of 91 neurons recorded by Huang et al. [13], containing 91 2-4 minute recordings, each from a different neuron. We narrowed our analysis to data from the three triple-transgenic mouse lines (out of the four presented in [13]) that were used by the Allen Brain Observatory. Two of the analyzed lines expressed the fast version of GCaMP6 (GCaMP6f) in L2/3 pyramidal neurons under the Emx1 and under the Cux2-promoter, respectively (abbreviated as Emx1-f and Cux2-f, respectively). The third line expressed the slow version of GCaMP6 (GCaMP6s) under the Emx1-promoter (abbreviated as Emx1-s). We used a numerical routine (described below) to select a subset of recordings for further analysis, in some cases including more than one recording per neuron (Tab.1). Our final data set was 50 recordings from 35 neurons across 3 transgenic mouse lines.

**Table 1.**
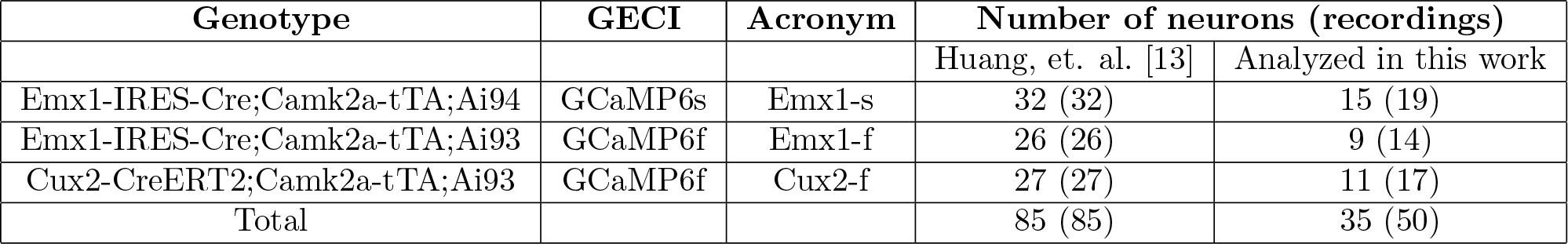
Composition of the calibration data set. Summary of the number of neurons, and individual recordings, broken down by genotype, genetically-encoded calcium indicator (GECI), and acronym used in the main text. See also QC-funnel diagram in Fig. 3D.

| Genotype | GECI | Acronym | Number of neurons (recordings) |  |
| --- | --- | --- | --- | --- |
|  |  |  | Huang, et. al. [13] | Analyzed in this work |
| Emx1-IRES-Cre;Camk2a-tTA;Ai94 | GCaMP6s | Emx1-s | 32 (32) | 15 (19) |
| Emx1-IRES-Cre;Camk2a-tTA;Ai93 | GCaMP6f | Emx1-f | 26 (26) | 9 (14) |
| Cux2-CreERT2;Camk2a-tTA;Ai93 | GCaMP6f | Cux2-f | 27 (27) | 11 (17) |
| Total |  |  | 85 (85) | 35 (50) |

### Electrophysiological data

#### Data preprocessing

The raw cell-attached recordings were baseline-corrected by subtracting a piece-wise polynomial fit (see S1 for details). Action potentials were detected as peaks of the baseline-corrected voltage traces, which exceeded the Quiroga threshold 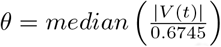 (an estimator of the background noise widely used in electrophysiology [15]) by a factor of 10 (see Fig. S1 for details).

#### Quality control

In order to quality-control (QC) the electrophysiology data, we developed a semi-automated procedure. In the absence of a set of universally agreed-upon metrics as to what constitutes a ‘good’ cell-attached recording, we created a ‘reference data set’ with the help of experienced human annotators (a group comprised of the authors and additional scientists listed in the acknowledgements). These experts crowd-sourced the data by reviewing a small subset of it, with each expert instructed to nominate any recording they would subjectively rate as ‘high-quality’, and to veto any recording nominated by others if they deemed it of questionable quality. This procedure resulted in a reference data set containing 45 vetted cell-attached recordings.

Second, we compiled a comprehensive set of quality-associated statistics, described in detail in Subsection Quality control metrics for cell-attached electrophysiology data. For each metric, a distribution was computed on the reference data set to define an acceptable range expected of high quality data. The metrics were subsequently computed on all of the available cell-attached data. A recording would pass QC if and only if for all metrics it fell within the range spanned by the reference data set. Fig. 1 A shows the distribution of the signal-to-noise-ratio (SNR) metric (defined as the ratio between the median spike amplitude and the Quiroga threshold *θ*) across the reference data set as well as across all of the data, and illustrates our approach to filtering data by quality. As shown in Fig. 1 B, some of the metrics were significantly more selective than others. Fig. 1 C, D show an example recording that passed QC and (Fig.1 E, F) one that failed QC. Overall 159 recordings (out of 553 with > 3 action potentials) passed this stage of the QC.

**Figure 1.**
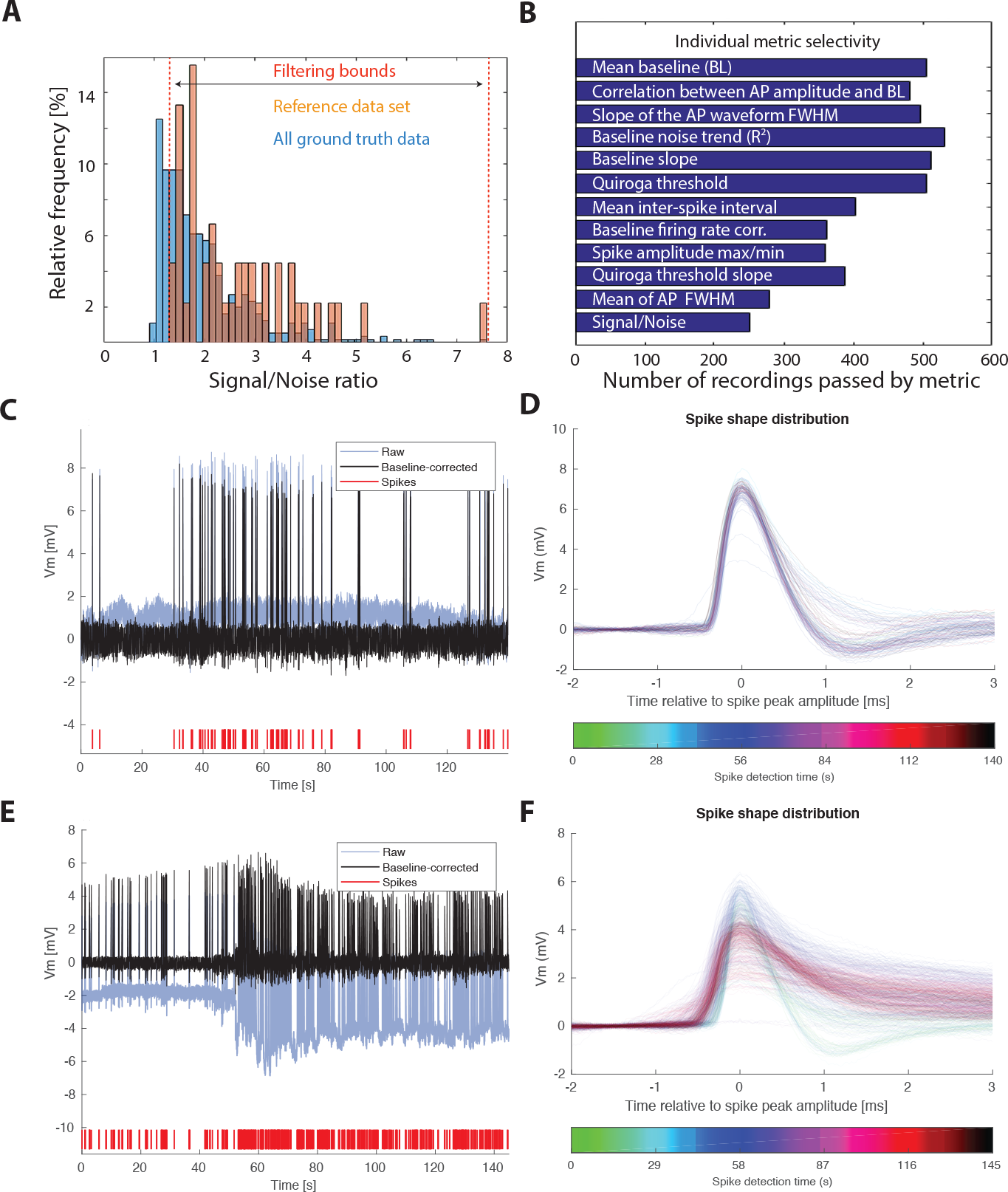
Quality control of electrophysiology data. **A** Distributions of the signal-to-noise-ratio across recordings in the reference data set (brown) and across all data (blue). **B** Selectivity for the 12 most selective QC metrics. See Appendix (Quality control metrics for cell-attached electrophysiology data) for a complete list of the employed metrics. **C** Example electrophysiology data that passed QC: overview of raw data (blue), data post baseline subtraction (black), and detected spikes (red). **D** Example cell-attached recording that passed QC: spike shapes; the color encodes the time during the recording when the spike occurred and facilitates detection of systematic drift or degradation. **E, F** Same as C, D but for a recording that failed QC due to a too high prevalence of spike shape broadening and variability.

### Detection and characterization of Prolonged Depolarization Events (PDEs)

We detected PDEs using an *ad hoc* thresholding approach in 50 ms intervals: the voltage trace was classified as free from PDEs if and only if 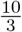 Quiroga threshold (which is 1/3 of the threshold we used for spike detection) was exceeded for ≤ 5*ms* per detected action potential. Otherwise, the interval was classified as likely containing a PDE. After identifying traces as containing such events we characterized them by amplitude (maximum amplitude above detection threshold) and duration (time the membrane potential remains above detection threshold past the peak of the last detected action potential).

### Two-photon imaging data

#### Data preprocessing

First, the two-photon movie was motion corrected using a custom algorithm, which implemented a translational correction based on cross-correlation in the Fourier domain (for additional details, please see Suppl. Section: Custom motion correction algorithm).

The movie was then sub-sampled by a factor of 4 in space, and a factor of 5 in time, to match the spatio-temporal sampling, and the approximate number of pixels per soma used in the Allen Brain Observatory [10]. To assess the effect of decimation on subsequent processing, 20 different approaches were tried in parallel (decimation starting with the 1st, 2nd, … 4th pixel × 1st, 2nd, … 5th frame, respectively). 4 × 5 internally identical blocks, one block for each decimation strategy, were tiled for a total of 400 almost identical regions of interest (ROI) per recording. These recordings were processed through the Brain Observatory image processing pipeline [10], which included region of interest (ROI) segmentation, filtering of the segmentation results by morphological characteristics, demixing (of overlapping somata), neuropil subtraction, and Δ*F/F*_0_-computation. Fig. 2Ai shows segmentation results, which are consistent across our family of sub-samplings, while Fig. 2Aii illustrates a case where the segmentation algorithm found more objects than just the patched neuron in the field of view. An example of neuropil subtraction is illustrated in Fig. 2 Bi.

**Figure 2.**
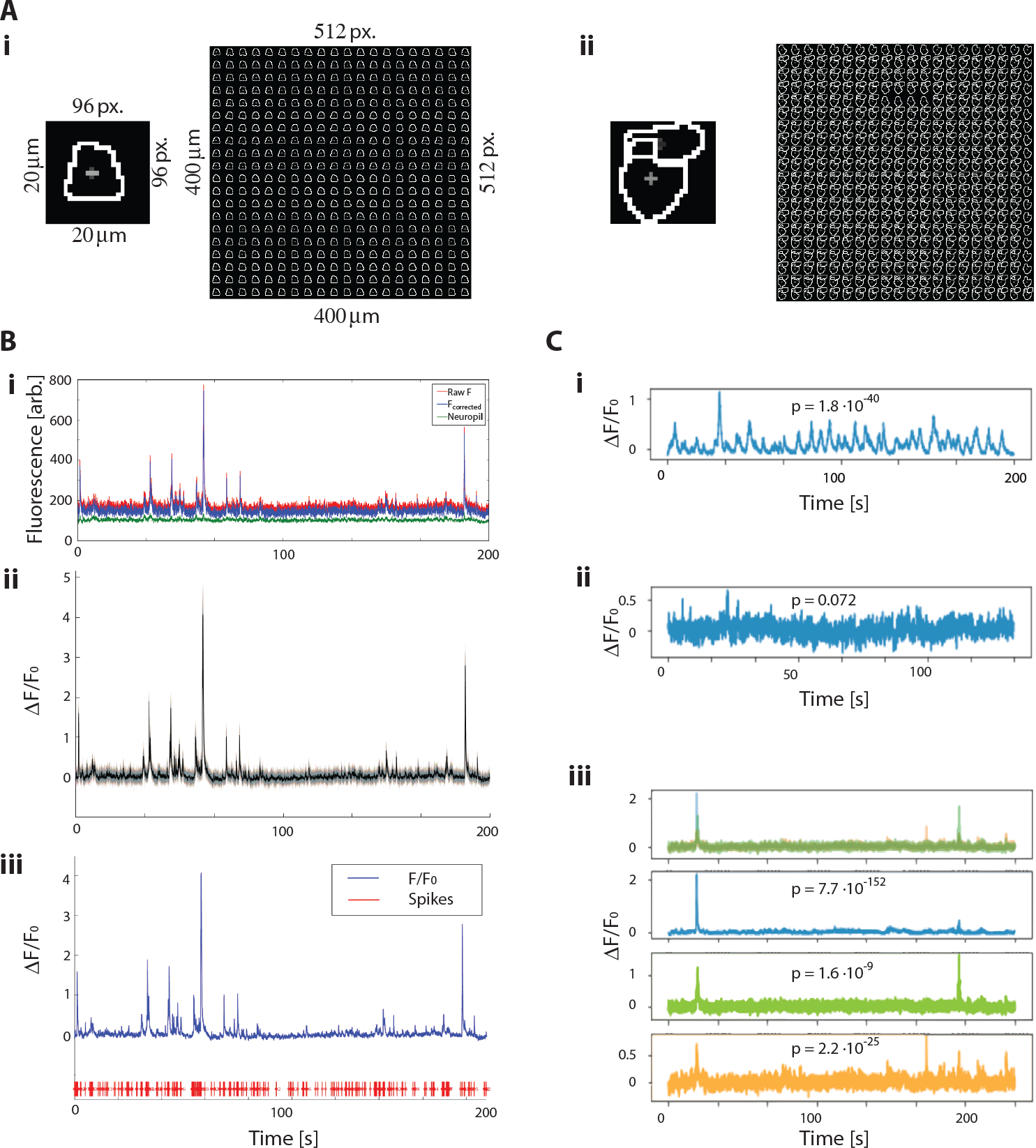
Quality control of two-photon imaging data. **A** Segmented regions of interest (ROIs) pre- and post-decimation and tiling; field-of view sizes are indicated before and after spatial decimation, respectively, i: Example ROI in which only segmented structures correspond to patched cell; ii: Some of the identified ROIs did not correspond to the patched cell. **B** Example of a high-quality recording that passed all QC criteria. i: Neuropil subtraction, ii: Self-similar family of fluorescence traces from 400 ROIs, median shown in black, iii: High quality fluorescence trace (blue) with measured spikes (red). Note many spikes do not appear to cause a significant change in fluorescent signal. **C** Examples of additional QC. i: Single cluster of traces that is significantly different from noise (passed KS-test), ii: Single cluster of traces that is not significantly different from noise (failed KS-test, as *p* > 0.05), iii: Color-coded overlay (top row) of three cluster medians (individual cluster medians shown in blue, green, orange), which all show different signals of likely neural origin (all three passed KS-test; p-values shown). The sum of all cluster medians had the highest correlation with the firing rate.

#### Automated quality control

In cases where only the patched cell was found during segmentation, trace extraction would yield a family of 400 self-similar traces (see Figs. 2 Bii and 2 Ci). To catch cases where segmentation yielded additional objects that were not part of the patched neuron of interest (see Fig. 2 Aii), additional QC steps were required. The traces were first clustered using DBSCAN [16, 17], and each cluster median was compared against white noise of the same mean and standard deviation (KS-test), and rejected as artifact if it was not significantly *p* < 0.05 different from Gaussian white noise of matched mean and variance, e.g. Fig. 2 Cii. In cases where multiple clusters were significantly different from noise (Fig. 2 Ciii), this was either due to multiple neurons being present in the field-of-view, or due to residual motion artifacts resulting in multiple translated copies of the same ROI. To disambiguate these two possibilities, the top three clusters were combinatorially merged, i.e. sums were computed for all six possible combinations (sampled without replacement) of (up to) three most distinct cluster medians, and the combination most significantly correlated with the measured electrophysiological spike train was picked. Correlation significance was determined by building a null distribution of correlations between the cluster medians and 1000 random Poisson trains with a rate matching that of the recorded spike train. If there was no more significant correlation between any cluster median (or sum thereof) and the measured spike train than the 0.5-th percentile of the null distribution (i.e. *p* > 0.005), the recording was failed. In total, 134 recordings passed this stage of the QC.

#### Manual QC refinement

This automated QC described is insensitive to very rare artifacts where recording quality started out high but degraded part-way through the experiment. Fig. 3A exemplifies a case where the cell was perforated half-way through the experiment, and the initially excellent correspondence between electrophysiology and optophysiology was lost. Nine such obvious artifacts were identified and the recordings containing them were eliminated from the analysis. Furthermore, since the data set was to be used for calibration, we wanted to avoid including recordings that showed artifacts expected only in simultaneous recordings but not in regular calcium imaging data, such as intermittent loss of the cell-attached patch (one possible explanation for ample activity clearly present in the fluorescence data but lacking of spiking correlates). We manually eliminated 33 candidate recordings that exhibited suspicious non-stationarities; an example is shown in Fig. 3B.

**Figure 3.**
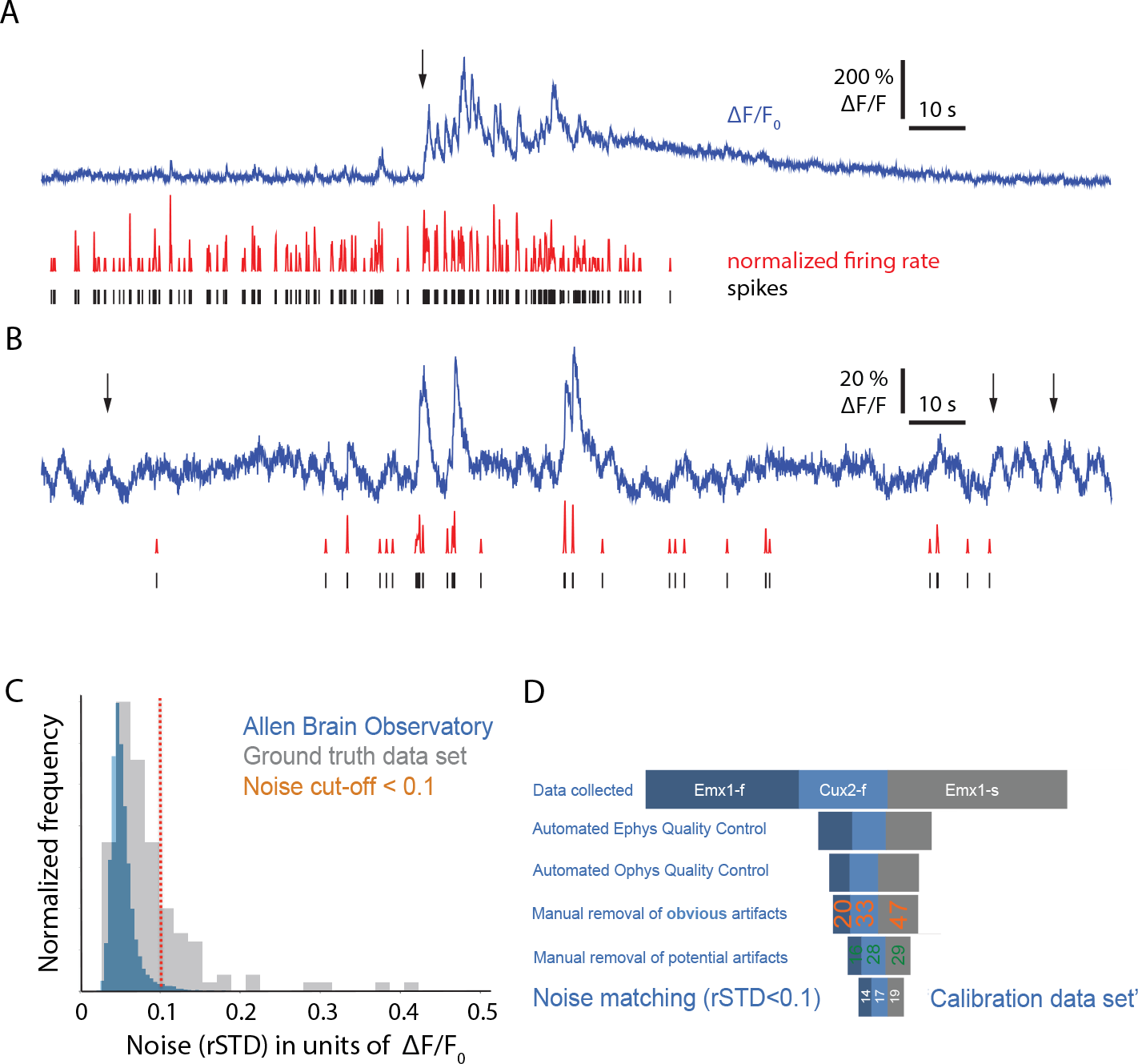
Summary of the quality control process. **A** Example of clear artifact removed during manual curation: the plot clearly shows a catastrophic event (indicated by black arrow around 70s) where the fluorescence drastically increases, but the cell ceases all electrical activity in under one minute after a brief initial period of high-frequency firing. In this example, the membrane was perforated, and extracellular electrolyte came in contact with the cytosol. **B** Example of likely artifact — recording was flagged and excluded during manual curation: significant fluorescence changes in the absence of electrical activity (indicated by black arrows around 8s, 112s, and 120s, respectively) most likely indicates a pathological state of the neuron or an artifact. **C** Distribution of the robust standard deviation of the noise across all of the Allen Brain Observatory recordings of 60,000 cells (blue), and across all artifact-free simultaneous recordings, respectively (gray). Most of the Brain Observatory data set featured a noise level below 10% Δ*F/F*_0_. **D** Visual summary of the entire quality control process (’QC-funnel’) that lead to our final ‘Calibration data set’ (at the bottom) of 50 recordings from 35 individual cells.

#### Filtering by noise profile

We computed the ‘robust standard deviation’ (a median-based method with outlier-removal) of the noise *rSTD* [10] for all remaining recordings, and only kept recordings where *rSTD* < 0.1, as is the case in the bulk of Allen Brain Observatory fluorescence data (Fig. 3C). Including this filtering step, the entire QC process yielded a grand total of 50 recordings. 13 of those recordings were of genotypes other than the three genotypes of interest in the context of the Brain Observatory. A schematic overview of the entire QC process is depicted in Fig. 3D. All further analysis is restricted to the 50 recordings summarized in Tab. 1.

#### Event detection

We used a state-of-the-art *l*_0_-regularized spike extraction algorithm [18] to detect the timing and the amplitude of events in the fluorescence portion of the data. An example trace post-QC and *l*_0_-event detection is shown in Suppl. Fig. S5 for each analyzed genotype. For comparison, we have also included events inferred using the biophysically inspired (but significantly slower) non-linear model MLSpike [19].

## SPICE simulations of different patching edge cases

We have modeled the cell-attached patching configuration via the lumped-element equivalent circuit model (Suppl. Fig. S7) in NI Multisim Live (SPICE): https://www.multisim.com/content/L42fQKhbiikEGUYdnb8kjn/cell-attached-recordings/. We modeled a spherical neuron, 10 *μ*m in diameter, with a patched area of *πμm*^2^, that is 1% of the total membrane area (100 *πμm*^2^). Approximated as a circle of ≈1 *μm* radius, this corresponds to a pipette tip 2*μm* in (inner) diameter, as used in [13]. Such a pipette has a tip resistance (*PipetteR*) of ≈ 5*M*Ω, implying an access resistivity (through the electrolyte) of *ρ*_*a*_ ≈ 15.7*M*Ω *μm*^2^, and an access resistance through a potential perforation of 1% of the patch area of *AccessR* = 500*M*Ω. We computed the capacitance *C* of the patched membrane and of the rest of the cell as *C* = *c* · *A*, where *c* denotes the respective membrane’s capacitance per unit area, and *A* represents the respective membrane’s area. We computed the resistance *R* of the patched membrane and of the rest of the cell as 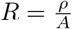, where *ρ* denotes the membrane’s resistivity. The values we used for membrane resistivity (*ρ*_*m*_ ≈ 51.8*G*Ω · *μm*^2^) and capacitance per unit area 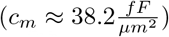 are consistent with values reported in the literature for mouse pyramidal neurons. The voltage source across the membrane is assumed to consist of an action potential followed by a prolonged depolarization event, both of which are approximated by voltage chirps connected in series. The voltage chirp parameters are summarised in Tab. 2.

**Table 2.** Voltage chirp parameters for simulated action potential with subsequent depolarization.

| Parameter | Action potential | Depolarisation |
| --- | --- | --- |
| Initial amplitude | 150 mV | 80 mV |
| Initial frequency | 500 Hz | 20 Hz |
| Final amplitude | 1 mV | 0.01 mV |
| Final frequency | 10 mHz | 100 mHz |
| Duration | 2 msl | 50 ms |
| Delay | 10 ms | 10 ms |
| Voltage Offset | -70 mV | 0 mV |

When the seal is poor, and the seal resistance is lower than the pipette resistance (e.g. ¡ 5MΩ), we see the ‘extracellular’ phase relationship, i.e. the patch pipette records approximately the derivative of the membrane potential, and the amplitude is in the hundreds of *μ*V, as one would expect for an extracellular recording. The minimum resistance at which ‘preparations for data collection began’ (Huang et al) was 20 MΩ. At this seal resistance, the AP amplitudes would already be in the mV range, and the phase of the recorded voltage would indicate an intermediate state between the polarity of an extracellular recording and that of an intracellular recording (albeit it still looks more extracellular and PDE is mostly high-pass filtered).

## Results

Our primary goal was to establish a quantitative link between the spiking activity of neurons and the calcium activity under common, desirable imaging conditions, namely using a large field of view (several hundred micrometers containing hundreds of neurons). We chose the openly available Allen Brain Observatory data set as our reference for desirable large field of view imaging conditions. We acquired images with a small field of view [13] and downsampled the images to match the increased noise of large field-of-view images, using the Allen Brain Observatory as a reference for the expected noise profile (see Methods). To obtain accurate spike times, we used cell-attached recordings.

In most studies, analysis is performed on a subset of recordings selected from a larger dataset, the intention of the selection process being to exclude recordings with inadequate signal-to-noise ratio or in which the neuron was perturbed by the recording pipette. Previous authors have selected recordings manually. Instead, we implemented an algorithm to select recordings, reducing subjectivity in the sub-selection process and enhancing reproducibility.

Possible perturbations that generate undesirable measurements include changes in cytoplasmic calcium concentration or in spiking activity, and can arise from several mechanisms. The most common mechanism being mechanical damage to the plasma membrane. If there is a perforation on the inside the patched portion of the membrane, the pipette has access to intracellular space, and the recording is no longer ‘cell-attached’ but a ‘whole-cell’ recording. Bringing the cytoplasm in direct contact with extracellular solution from the pipette would permit entry of extracellular calcium into the neuron, elevating the intracellular calcium concentration.

The relationship between the electrical characteristics of the patched cell and the observed voltage waveform can be complex. To assess the types of waveform we should expect in our data, we considered how seal resistance (resistance from the pipette tip to the extracellular space through the seal formed by the cell membrane against the pipette tip) affects electrical recordings. We found through lumped-element SPICE simulations (see Suppl. Fig. S7) that the seal resistance determines whether cell-attached recordings (intact patch membrane) look more like attenuated whole-cell recordings (low access resistance when seal resistance is high), or more like large-amplitude extracellular recordings (very high access resistance and low seal resistance), i.e. the first derivative of a whole-cell recording. While only selecting recordings that clearly look extracellular would effectively exclude whole-cell recordings, it would also preclude further analysis of cell-attached recordings with high seal resistances, likely those with the highest signal-to-noise ratios. We included a red indicator (Alexa 594) in the recording pipette and excluded from further analysis all recordings with red indicator in the soma. [13].

Our algorithm was based on a series of numerical criteria that were applied to the electrophysiology traces and calcium images of every recording. Only recordings that passed every criterion were subjected to further analysis. For the electrophysiological recordings, these criteria were established by first having a round of experts annotate traces as ‘high-quality’. These traces were used to establish a distribution of metrics that defined the class of ‘high-quality’ recordings and data was included if its values for these metrics fell within the range of that for the annotated ‘high-quality’ recordings. This process of quality control reduces the variability introduced by individual annotators and puts quantitative measures on a ‘crowd-sourcing’ of expertise in electrophysiology. Further information on the algorithm for both optical and electrical recordings is provided in the Methods. The full data set and selection code are supplied, permitting reproduction of our results and adaptation of our routines to additional data sets.

### Expected probability of event detection and expected event magnitudes

The primary question of interest in establishing a quantitative link between the spiking activity of neurons and the calcium activity is to determine whether and how well one can identify spiking events from fluorescence traces. We expected the probability of detecting a spike to depend on the firing rate, the estimation of which is affected by the length of the window over which we count spikes. We measured the accuracy of event detection with the *l*_0_ penalized routine of [18]. Since the true spike train is known for these data, we were able to directly compute the probability of detecting an event, as well as the expected event magnitude, as a function of the true number of spikes observed in a detection window of a given duration. The results of this analysis are shown in Fig. 5. The chosen detection windows represent typical time scales over which averaged activity was defined in [10] using data from the Allen Brain Observatory.

**Figure 4.**
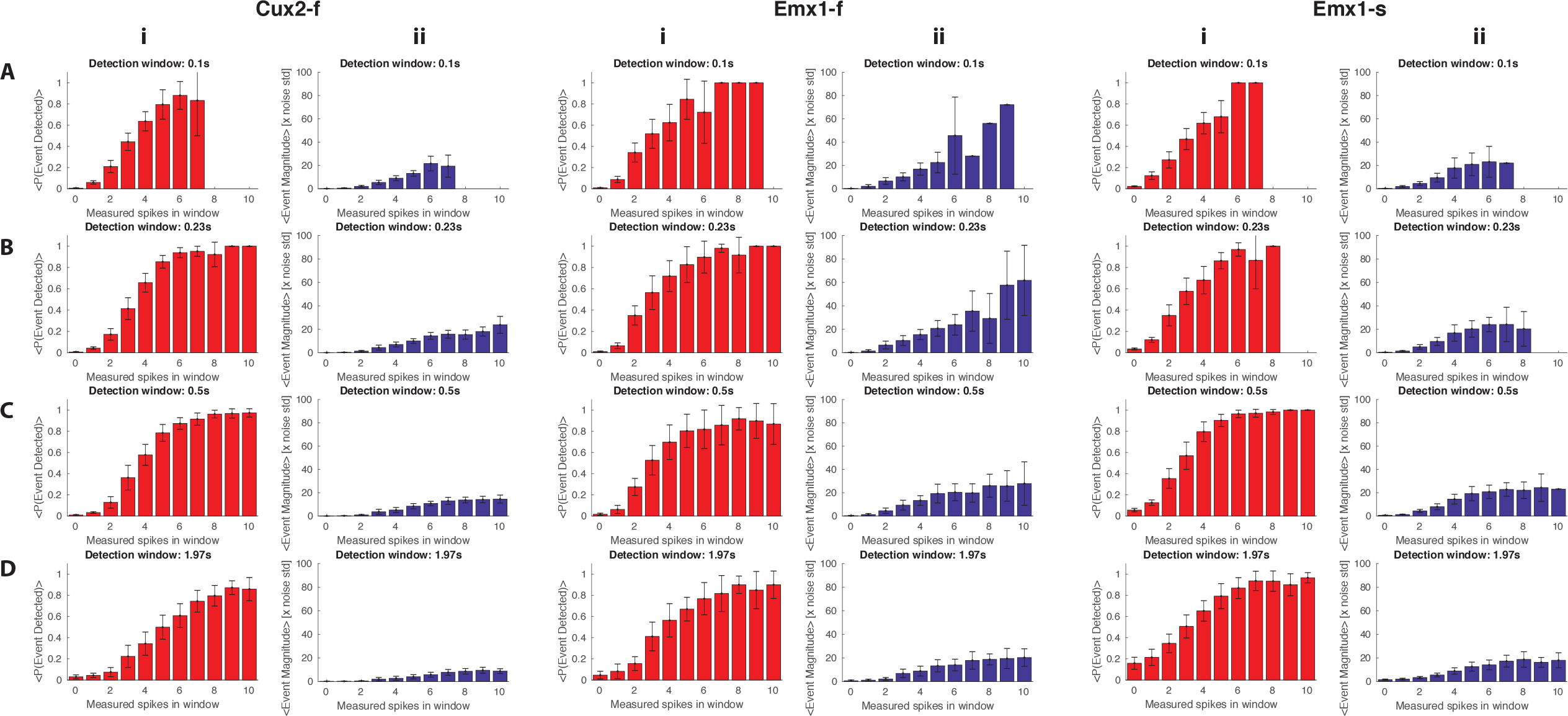
Expected probability of event detection and expected event magnitude. **A** i: Expected event detection probability (red) and ii: magnitude (blue) for a detection time window of 0.1s. The genotype is indicated in the column title. All error bars denote 2x standard error of the mean. **B, C, D** As in A but for a detection time window of 0.23, 0.5 and 1.97s, respectively.

**Figure 5.**
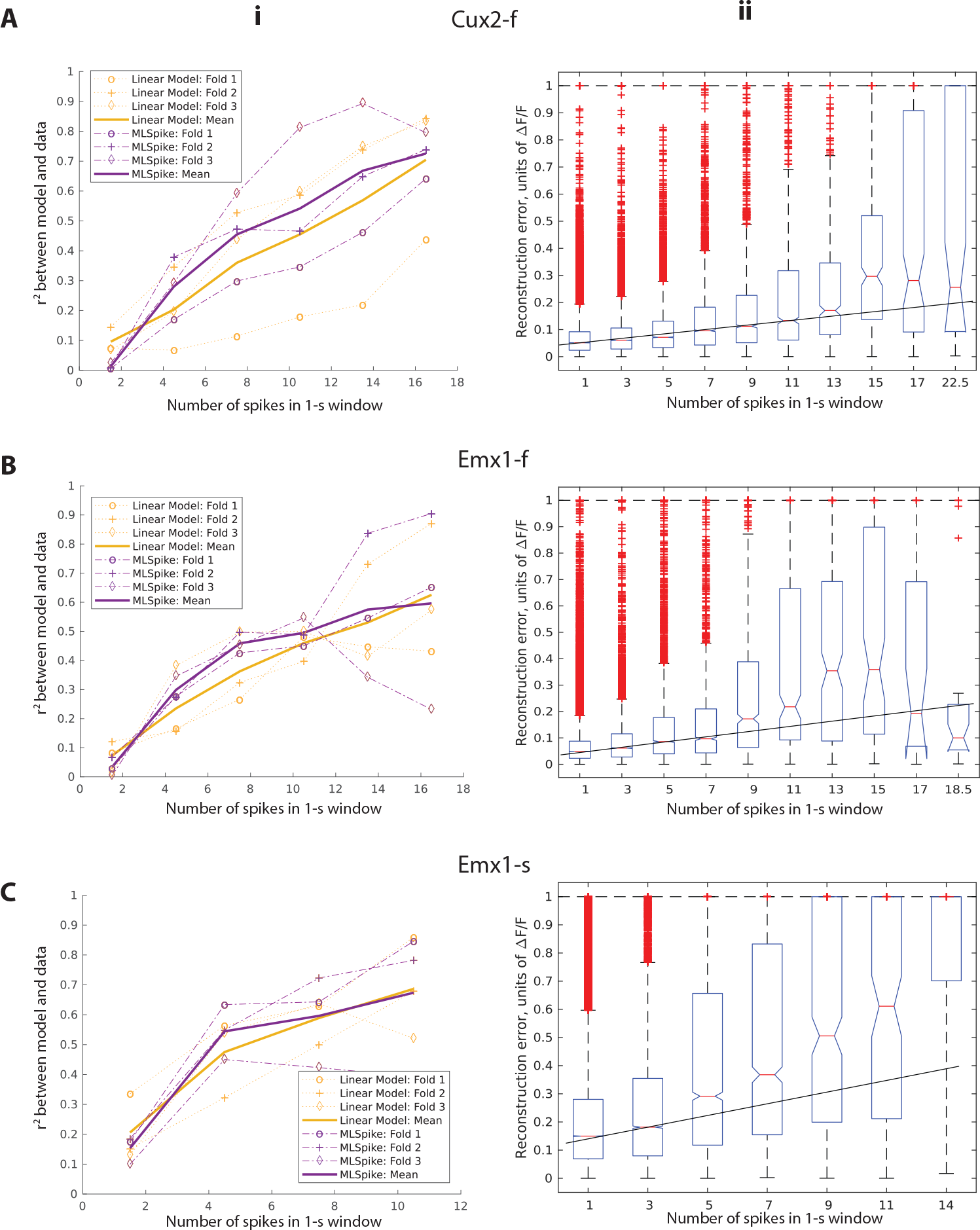
Detailed analysis and model comparison. **A** Analysis results for Cux2-f. i: Model performance as a function of firing rate for two different models: a simple linear model, which assumes that the fluorescence is the convolution of the spike train with a linear kernel (see Supplementary Fig. S2), and the bio-physically inspired non-linear model MLSpike [19]. MLSpike performed slightly better than the linear model on average but not on all folds (disjoint, consecutive subsets of data). Both models’ performance was 3-fold cross-validated using held-out data, see Suppl. Fig. S3 for more details. ii: Reconstruction error for the linear model as a function of the firing rate. Cux2-f shows a greater range of linearity (up to 11 Hz) than Emx1-f (up to 7 Hz) and Emx1-s (up to 3 Hz). **B, C** As in A but for Emx1-f and Emx1-s, respectively. MLSpike does not perform significantly better than the linear, convolutional model.

As expected, the probability of detecting spiking events was greater for multi-spike events than for single spikes, in all 3 mouse lines and with all detection window lengths. Almost all events containing 5 or more spikes were detected with GCaMP6f and GCaMP6s, at all detection window lengths. In contrast, the mean probability of spike detection was < 0.1 for single-spike events in both GCaMP6f lines and greater than 0.1 with GCaMP6s only with the 2s detection window. The detection of 2-4 spike events differed across mouse lines and detection windows (Fig. 5). Importantly, the false positive rate (the probability of detecting an event in the absence of spikes) was low (≪ 0.1) in all mouse lines for all but the longest window length. In longer time windows, a fixed spike count may result in a lower instantaneous firing rate, leading to a decrease in the average detection probability. For similar reasons, expected event magnitudes tended to decrease for longer event detection windows. We conclude that most of the events detected in the Allen Brain Observatory likely result from multiple spikes. Moreover, our results suggest that as many as 4 spike events are commonly overlooked in population imaging studies using GCaMP6f and GCamP6s.

Each spike inference or event detection algorithm has a unique set of strengths and weaknesses and this raises the question of whether our results are specific to the method of [18]. We tested the robustness of the result shown in Fig. 5 to the choice of specific algorithm used to extract events from the fluorescence time series by performing spike inference with MLSpike, shown in Fig. S4. Given the very different nature of these two routines (one based on a simple dynamical model of calcium, the other inspired by biophysics), we expect these results to hold for any general spike inference method. This conclusion is supported by the results of the next section.

### Modeling calcium-associated fluorescence from spikes

The low probability of detecting a calcium event for low spike rates raises the question of whether we can predict the calcium response given the spiking activity. If one aims to reconstruct a spike train from calcium activity, then the calcium activity needs to contain a consistent predictable signal derivable from the spikes.

To test this, we trained models of calcium response using two methods, a relatively simple linear convolutional model and MLSpike, a biophysically-inspired model that explicitly models the cooperative binding of calcium to GCaMP [19]. We use these models to predict fluorescence changes from spike times, determining prediction accuracy for events containing different numbers of spikes. We expect that in general a model will not be able to accurately identify spiking events from fluorescence if one cannot accurately predict fluorescence from spike times. Hence these models provide some insight into which types of spiking event are amenable to detection.

Figs 4A-C show the performance of these models (characterized by *r*^2^ between the predicted and true calcium signal) as a function of the number of spikes in a one second window. The calcium is never perfectly predictable from the spike rate, as we would expect, but the *r*^2^ values for low spiking events are particularly low 0.1, indicating that low spike rate periods produce no appreciable, consistent calcium response. For the purpose of model comparison, each model’s parameters were trained on two thirds of the total available data for each genotype. Testing was performed on the held-out third, and boot-strapped, yielding three folds, i.e. data used for testing was never used for parameter estimation. (Suppl. Figs. S3A-C i, ii show how well the models predicted the measured fluorescence data on the testing set for each fold.) Both models explain very little variance at low firing rate, and steadily improve for increasingly higher firing rates, at which more of the fluorescence fluctuations were driven by underlying action potentials. The improvement is more rapid for GCaMP6s (than for GCaMP6f), which offers larger and longer-lasting fluorescence responses per spike (Suppl. Figs. S2 C). Notably, there is no significant performance difference between the simple linear model, and the more complex non-linear model (MLSpike).

The linear convolution model assumes that the fluorescence can be expressed as the convolution of the underlying spike train with a linear kernel. Suppl. Figs. S2 A-C show optimal linear kernels (See Supplement for details of linear kernel extraction) for the three investigated genotypes. It is worth noting that one should resist the temptation to interpret these kernels too literally as the physical fluorescence responses to single, isolated spikes. Rather, they represent average responses that under a linear model, minimize the mean squared error. True single-spike responses, when detectable, would decay faster [9]. A comparison of panels Suppl. Figs. S2 A-C shows that the shape of the optimal kernel is largely determined by the properties of the GECI (i.e. GCaMP6f versus GCaMP6s), rather than by the driving promoter (Emx1 versus Cux2). We determined the half life of the linear kernel via single-exponential fit (starting at kernel maximum). The amplitude was largest (13.0*±*4.9%), and the half-life was longest 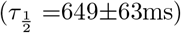, for Emx1-s, second-largest (8.6*±*1.1%), and second-longest 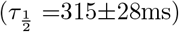 for Emx1-f, and smallest (6.5*±*0.6%) and shortest 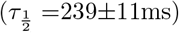 for Cux2-f. The variability of the kernel amplitude and decay time also decreased in then same order: Emx1-s *≫* Emx1-f > Cux2-f. The reporting kinetics is consistent with Chen et al. [9]: the half-lives we extracted lie in-between those reported in [9] for single action potentials and those reported for 10 action potentials.

If the linear convolutional model described the data well, we would expect the reconstruction error to be a linear function of the firing rate. Indeed, this is the case for low firing rates (Figs. 4 A-C ii), but the linear relationship progressively breaks down at higher firing rates as the GECI become more saturated with Ca^2+^. Notably, this loss of linearity occurs at lower firing rates for GCaMPs (>3Hz) than for GCaMPf (>7-11Hz).

The agreement of both a simple linear, convolutional model and the detailed biophysically based MLSpike demonstrate that there is very little predictable relationship between calcium and spiking activity at low firing rates, indicating that other processes are determining much of the observed calcium dynamics.

### Prolonged depolarization events associated with elevated cytoplasmic calcium

Inference of the timing of action potentials from calcium indicator fluorescence generally operates on the premise that action potentials are the dominant source of calcium influx. In some instances, spikes do not appear to be the dominant source of calcium, a fact reflected in the relatively poor performance of the models of calcium from spiking activity that we tested. We observed that in a sub-set of neurons some action potentials were followed by prolonged periods of depolarization lasting from *≈*5 ms to >50 ms Fig. 6 A. We term these Prolonged Depolarization Events (PDEs) and asked to what extent these depolarizations were predictive of calcium activity.

**Figure 6.**
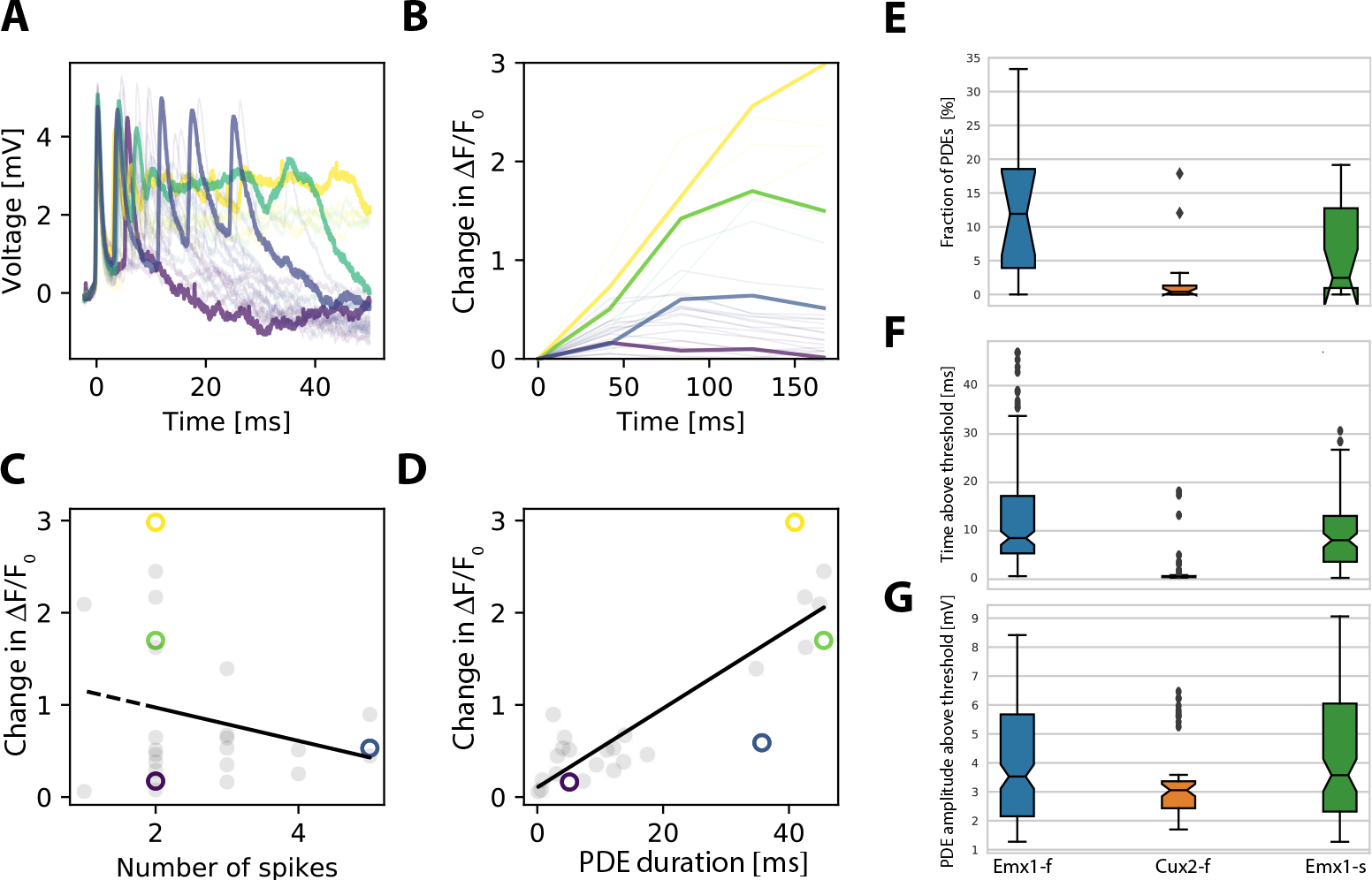
Prolonged Depolarization Events (PDEs) correspond to the largest fluorescence responses. **A** Four 50 ms snippets from the cell-attached voltage trace of a single Emx1-f neuron; the snippets contain 2-5 spikes within the 50 ms window and feature PDEs of variable duration. The color-code denotes corresponding elements between the four panels of this figure. **B** Change in Δ*F/F*_0_ over 150 ms following the electrophysiological event (color-coded to reflect response amplitude, and to match the corresponding voltage trace). **C** A scatter plot and linear regression of the number of spikes detected in the voltage snippet against the fluorescence response. **D** A scatter plot with linear regression of the PDE duration versus the changes in in Δ*F/F*_0_. Snippets corresponding to other spiking events and their corresponding data points in the other panels are greyed out for clarity of illustration. The greyed out traces, with additional detail, as well as data for other Cre lines, are shown in Suppl. Fig. S6. **E-G**: PDE prevalence (E), duration (F) and amplitude (G) for our three mouse lines.

We define PDEs as periods of time following action potentials (or bursts of action potentials) where, after the peak of the last action potential, the membrane potential exceeds a threshold (that is lower than the action potential detection threshold) and is maintained for a length of time that is longer than an action potential (≈ 2*ms*). The precise quantitative details of sub-threshold depolarisation detection and characterization (including choice of threshold) are outlined in Methods.

There is a relationship between the magnitude of the resulting fluorescence response and the duration of the PDE (Fig. 6 B). When PDEs are present, the number of spikes preceding a fluorescence transient tends to be a much weaker predictor of the transient’s magnitude (see regression of fluorescence response against the number of spikes in Fig. 6 C) than the PDE (see regression of fluorescence response against the PDE duration in Fig. 6 D), empirically defined as the time the membrane potential remains above threshold (see Suppl. Fig. S6 for additional details).

We observed significant differences across Cre lines in metrics of the PDE (Fig. 6E-G), namely the fraction of such events (Fig. 6E), the duration (Fig. 6F), and the amplitude (Fig. 6G). In 8/14 (57 %) recordings from the Emx1-f line, 6/19 (32 %) in Emx1-s and 3/17 (18 %) in the Cux2-f lines, significantly more of the variance in the fluorescence response could be explained by the presence and duration of prolonged PDEs that followed spikes than by the number of action potentials immediately preceding the fluorescence increases. Examples from all three Cre lines are presented in Suppl. Fig. S6.

In summary, our results provide evidence for physiological dynamics, namely PDEs, not associated with fast, voltage-gated Na^+^ channel-dependent action potentials but that are predictive of large somatic calcium transients in some Cre lines. The duration of PDEs, in recordings where they are present, accounts for a much larger fraction of the observed fluorescence changes than does spike count. As such, PDEs are an example of non-spiking dependent calcium dynamics which, unless taken into account and modeled explicitly, frustrate any attempts at recovering spiking activity from fluorescence measurements.

## Discussion

Calcium imaging is used to measure neural activity *in vivo*, often of hundreds or thousands of neurons across a large field of view, and is commonly expected to produce a proxy for action potentials or firing rates. However, the relationship between action potentials and fluorescence is non-linear and complex, necessitating calibration that is often performed with simultaneous imaging and cell-attached recording, at high frame rates and with the neuron of interest filling the field of view of the microscope to ensure the highest possible signal-to-noise ration; While these imaging conditions are optimal for indicator characterization or the calibration of single-cell studies they do not mimic typical population imaging experiments. Imaging with a larger field of view may result in a shorter pixel dwell time, poorer signal-to-noise ratio, and likely a reduced detectability of action potentials. Obtaining the map from measured fluorescence to spiking activity is of principal importance for this approach to systems neuroscience. Our aim here was to describe this map under common imaging conditions. As a reference for standard imaging, we used the Allen Brain Observatory [10] to serve as an exemplar of standardized, and quality-controlled population-scale somatic imaging studies for systems neuroscience.

For calibration, we obtained two photon calcium movies of neurons in three mouse lines with simultaneous cell-attached electrophysiological recording, as described in [13]. Most of the data was collected with a small field of view and at high imaging frame rates. We resampled the movies to match the larger field of view, lower sampling rate, and higher noise level of the data from the Allen Brain Observatory. The result approximated a ground truth data set for population scale recording conditions. Along with the original data from [13], this resampled dataset is publicly available.

Any experiment will have various sources of noise, systematic and otherwise, that must be identified and controlled. This is often accomplished by the manual selection of recordings for further analysis from the super-set of experiments. To reduce subjectivity and increase transparency and reproducibility, we adopted a quality control scheme that may be useful to other labs. We employed multiple experts to annotate traces, defining a core, high-quality data set. We then generated quantitative metrics for inclusion criteria based on this core data set. This approach converts the “wisdom of the crowd” into a quantitative and transparent algorithm for data inclusion.

There have been many attempts at deriving spikes from fluorescence data and it is recognized to be a challenging problem ([18–24] and others). Our results quantify the detectable calcium signal as deriving effectively from periods of relatively high instantaneous firing rate, highlighting an important bias in two-photon imaging, at least for our choice of population-scale field of view, indicator, mouse-lines, and *in vivo* conditions. Specifically, based on our results, observing a calcium event implies there are more than a few spikes (≤ 3) within a short window of time (100 − 500*ms*) (see Fig. 5). We expect these results to qualitatively hold for similar imaging conditions and commercially available instruments.

We chose to quantify the relationship of calcium to spiking activity by first converting the fluorescence traces to “calcium events” using a “spike” inference algorithm. Our results indicate that the calcium events identified via spike inference algorithms have a weak relationship to spikes at low firing rates. This is the primary reason we insist upon the language “calcium event” even when using a “spike” inference algorithm (this language was also adopted in [10]). We have shown this explicitly for two types of “spike” inference models. We expect these results to generalize to other approaches. First, the model of calcium dynamics used in the *l*_0_-penalized method of [18] is the same dynamical model found in other approaches to spike inference, even though it uses a different optimization scheme and penalty. Second, while one may wonder if the simplified calcium dynamics of the model in [18] (and others) is the culprit of our inability to derive spikes from fluorescence traces, the more biophysically realistic model of MLSpike fares no better (see Fig. S4). We therefore expect the set of commonly used approaches for spike inference to produce qualitatively similar results.

Conversely, we found that converting spikes to fluorescence traces was also problematic at low firing rates as we were unable to predict the calcium response reliably for low firing rates. Performance was similar for a simple model (linear convolution) and a biophysically more realistic model (MLSpike), suggesting that these results are general and not an artifact of model selection. Moreover, these results suggest that there is a fundamental problem in spike inference. If calcium cannot be consistently predicted from spiking events, then one cannot hope to recover the spikes from the recorded calcium.

The failure to convert spikes to fluorescence was not only the result of low signal-to-noise in our recordings. In some Cre lines we identified events in the voltage traces, which we termed here Prolonged Depolarization Events. PDE durations were strongly predictive of fluorescence, whereas the number of spikes immediately preceding or coincident with the fluorescence changes was not.

We observed prolonged subthreshold depolarizations (PDEs) associated with increases in somatic calcium concentration. The mechanisms underlying PDEs are unclear. Unlike previous calibration experiments, we examined neurons in transgenic mice. PDEs were most common in Emx1-IRES-Cre mouse line crosses, a line which is prone to epileptiform activity [25]. Extraction of spiking information from calcium imaging measurements in some transgenic mouse lines may require extension of spike inference models to incorporate additional sources of calcium with different characteristics from those evoked by action potentials.

In summary, we used the data set in [13] to calibrate the spike to calcium relationship under imaging conditions similar to those found in the Allen Brain Observatory [10]. Not only was the link between spike times and calcium transients weak at low firing rates, but calcium was difficult to predict from spike times. For some mouse lines, establishing the relationship between fluorescence and spikes may be impossible without more information than is typically available from population calcium imaging experiments.

## Acknowledgments

We would like to thank David McCormick, Sam Gale, and Corbett Bennett for their assistance with the curation of the reference data set for quality control of the cell-attached recordings. We wish to thank Gabe Murphy for helpful discussions. We wish to thank Carol Thompson and John Phillips for their help and support in managing this project. We further thank Mark T. Harnett for helpful discussions regarding afterdepolarizations. We wish to thank the Allen Institute founder, Paul G. Allen, for his vision, encouragement and support.

## Author Contributions

CK, MAB, HZ conceived the project, LL, UK, LH performed experiments. PL, LH analyzed data. RCR, CK, MAB, HZ, SEJdV, JW supervised the project. PL, MAB, and JW wrote the paper with input from LH, SEJdV, and CK.

## Competing Interests

The authors declare no competing financial interests.

## Data availability

The calibration data set will be available on the Allen Institute website: https://portal.brain-map.org/explore/circuits/oephys

## Source code availability

The source code used for QC and analysis will be available on GitHub, expressly free of support or warranty

## Supplementary Material

### Quality control metrics for cell-attached electrophysiology data

Metrics computed on continuous electrophysiological data:

- Median relative deviation of the membrane potential (MRDM), i.e. ratio between the median absolute deviation (MAD) and the median: 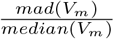
- Mean of the Baseline (BL)
- Baseline coefficient of variation: 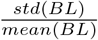
- Baseline noise mean, as approximated by the Quiroga threshold (QT) [15]
- Stability of the QT: 1000 10s-intervals were uniformly sampled from each recording, and the QT was computed on each sample. Quiroga Noise Stability (QNS) was defined as the coefficient of variation over the 1000 QT samples.
- *r*^2^ value obtained from a linear regression (Matlab’s `regression` function) of the 1000 QT samples against the start times of the 10s segments on which the QT was computed.
- Slope obtained from a linear regression (Matlab’s `regression` function) of the 1000 QT samples against the start times of the 10s segments on which the QT was computed.
- *r*^2^ value obtained from a linear regression (Matlab’s `regression` function) of the baseline against time.
- Slope obtained from a linear regression (Matlab’s `regression` function) of the baseline against time.
- Negativity: 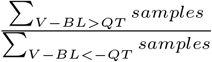

Metrics computed on the detected action potential time series (only recordings with > 3 action potentials, otherwise the recording automatically failed QC):

- Number of spikes
- Estimate of the maximum likelihood inter-spike interval via Matlab’s `lognfit` function.
- Mean spike amplitude
- Spike amplitude coefficient of variation
- Spike amplitude median relative deviation
- Relative spike amplitude range: 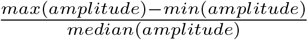
- Spike amplitude Max/Min ratio: 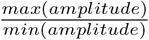
- Signal-to-noise ratio (SNR), defined as: 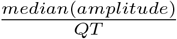

Metrics computed on the shape of the 2ms-long spike waveforms (±1ms around detected spike time) smoothed using Matlab’s `smooth` function with `sgolay` option:

- Mean Width at half the amplitude before the detected spike time (’left’ width-half-max, LWHM)
- Mean Width at half the amplitude after the detected spike time (’right’ width-half-max, RWHM)
- Mean Width at half the amplitude around the detected spike time (full-width-half-max, FWHM): *FWHM* = *LWHM* + *RWHM*
- Coefficient of Variation of the LWHM
- Coefficient of Variation of the RWHM
- Coefficient of Variation of the FWHM
- *r*^2^ value obtained from a linear regression (Matlab’s `regression` function) of spike amplitude against spike time.
- Slope obtained from a linear regression (Matlab’s `regression` function) of spike amplitude against spike time.
- *r*^2^ value obtained from a linear regression (Matlab’s `regression` function) of spike FWHM against spike time.
- Slope obtained from a linear regression (Matlab’s `regression` function) of spike FWHM against spike time.

Firing-rate based metrics (Firing rate was estimated by convolution of the spike train with a 1s-long box-car window using Matlab’s conv function):

- Mean firing rate (FR)
- Coefficient of Variation of the FR
- *r*^2^ value obtained from a linear regression (Matlab’s `regression` function) of firing rate against time.
- Slope obtained from a linear regression (Matlab’s `regression` function) of of firing rate against time.
- Pearson correlation (Matlab’s `corrcoef` function) between the baseline and the firing rate.
- Pearson correlation (Matlab’s `corrcoef` function) between the baseline at the time spikes were detected and the spike amplitudes.
- Pearson correlation (Matlab’s `corrcoef` function) between the baseline at the time spikes were detected and the spike FWHM.

## Custom motion correction algorithm

The Allen Brain Observatory pipeline contains a 2D correlation-based (2D FFT) motion correction step, which corrects for xy-translation when there is enough static structure in each frame to obtain a robust estimate of displacement. When recording from large fields of view containing hundreds of neurons, such structure is almost always present. In contrast, the two-photon data we analyzed in this work often only captured a single neuron (or, at most, a few neurons) in the field of view. Thus, whenever the neuron was inactive, the FFT phase would be dominated by noise, and our standard approach to motion correction had to be modified. The raw calcium movie, sampled at up to 169 samples/s, was sub-divided into sub-stacks, which were averaged and the averages aligned, such that there was always enough signal for meaningful motion-correction. The displacement vectors were then used to translate the individual frames comprising the sub-stacks, and the process was repeated with more, smaller sub-stacks (first 2, then 4, 8, etc sub-stacks down to 5 frames per sub-stack). With each iteration of alignment, progressively higher-frequency motion was corrected for.

### Linear kernel extraction

Linear convolution kernels were extracted by regularized linear regression (*L*_2_-penalty) using the `Tensorflow` back-end of `Keras` in Python 3.6. Snippets of the observed spike train, 256 samples in length, were used as input to a 1D convolutional layer with a rectifying activation function, and the network was trained to approximate the corresponding snippets of fluorescence, thus learning filters, which convolved with the spike train, approximated the measured fluorescence response.

---

**Figure S1.**
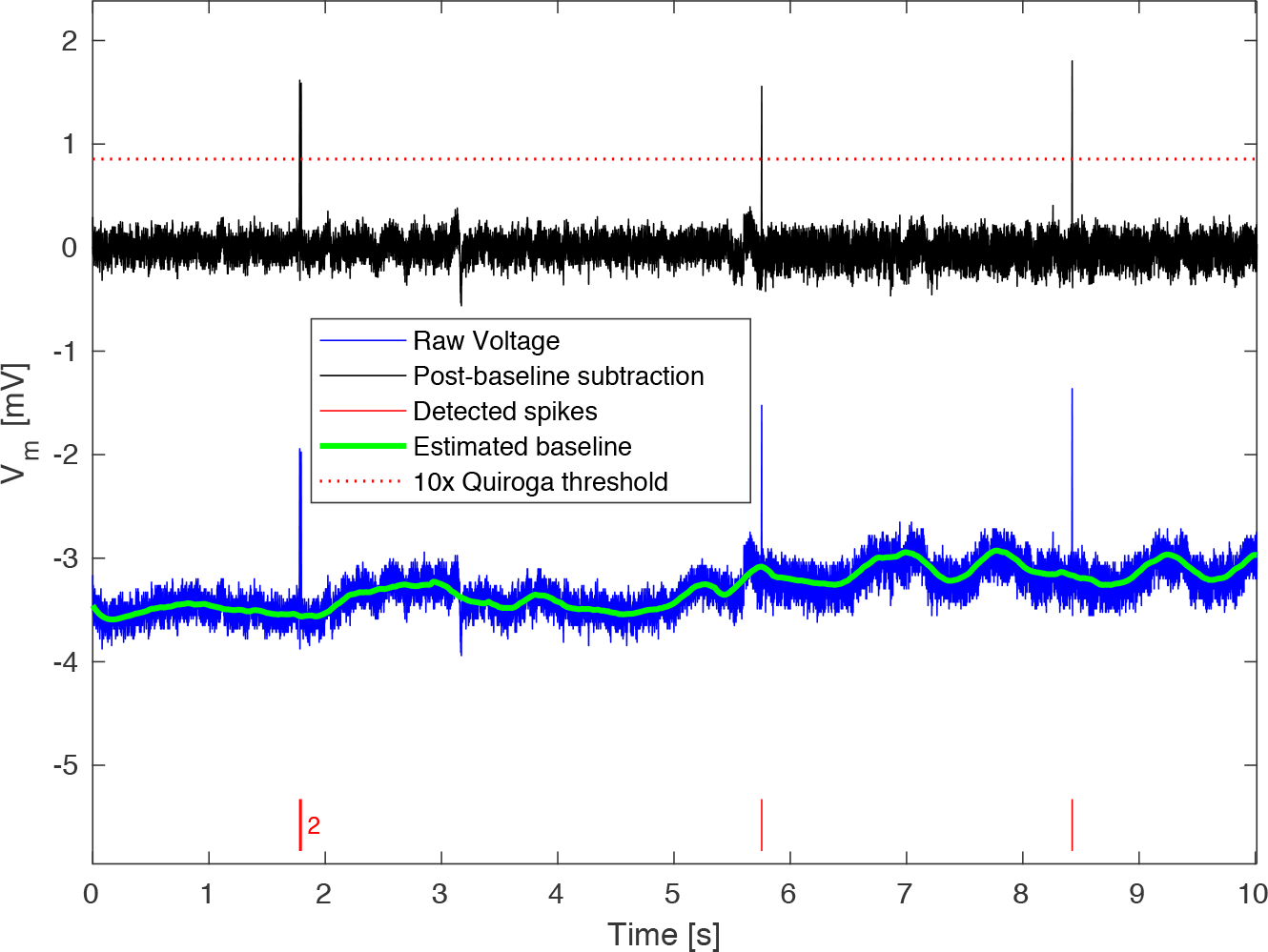
Preprocessing of electrophysiological data. Raw voltage traces (blue), (sampled at 40 kHz), commonly contained large baseline fluctuations that complicated spike detection. Prior to spike detection, such baseline fluctuations were removed by baseline subtraction. The baseline (green) was estimated by Savitzky-Golay (polynomial smoothing) filtering (3-rd order filter over a frame length of 20,001 samples using Matlab’s built-in `sgolayfilt`) of the raw voltage trace. In this example, four spikes (red raster) are detected when the baseline-corrected trace (black) crosses 10x the Quiroga threshold (red dotted line). Two of the spikes are separated by a very short inter-spike interval (raster ticks marked with red ‘2’).

**Figure S2.**
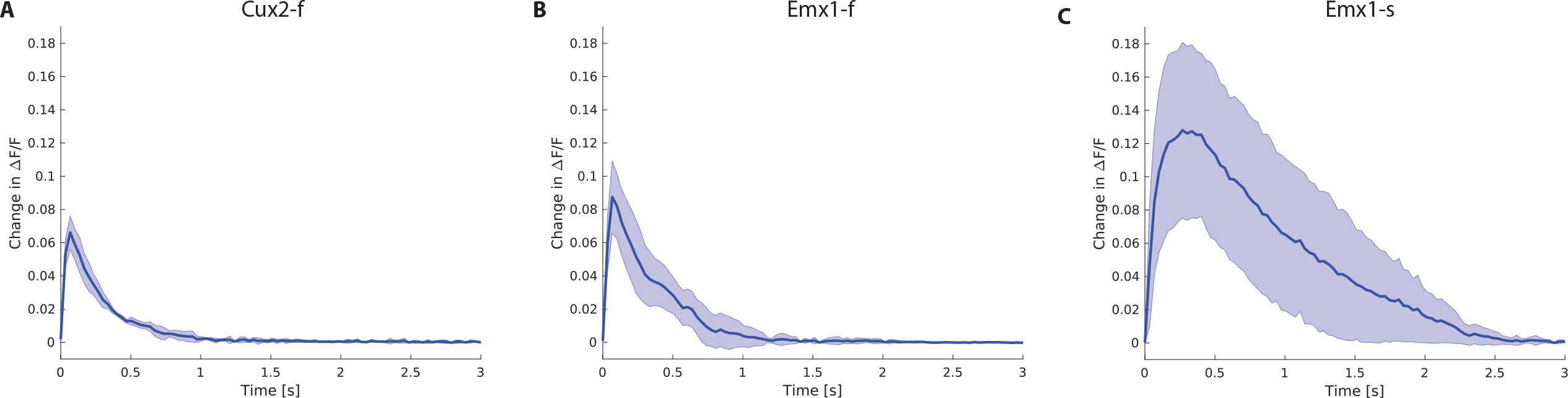
Extracted linear kernels. **A** Analysis results for Cux2-f. i: Optimal linear kernel. The error envelope is 2x the standard error of the mean across folds; **B, C** As in A but for Emx1-f, and Emx-s, respectively.

**Figure S3.**
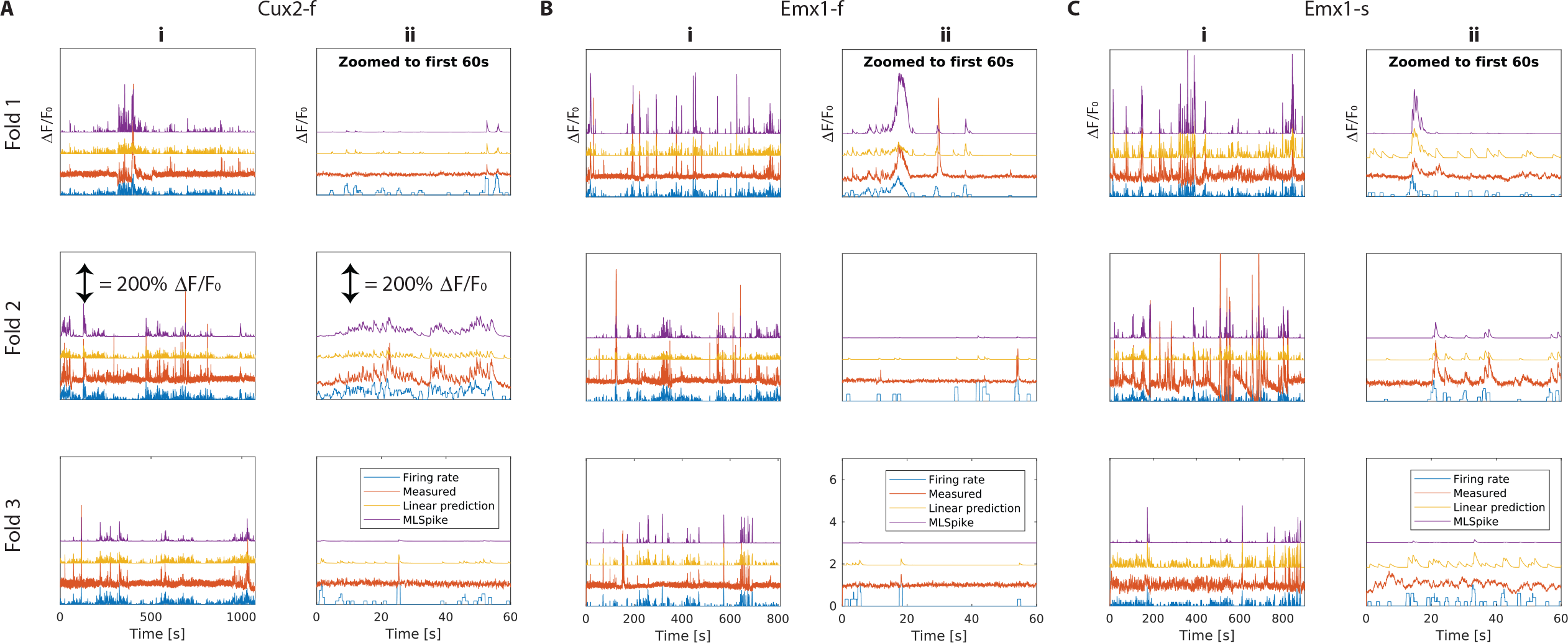
Detailed comparison of model performance across folds. **A** Analysis results for Cux2-f. i: the duration of the entire validation fold, ii: zoomed in on the first minute of the validation fold. Predictions by simple linear model (yellow) and by MLSpike (violet) superimposed on raw fluorescence (red) and an estimate of the firing rate (convolution of the spike train with a 1s-long box-car window; plotted in blue). MLSpike preforms marginally better on average than the naive linear model but it does not outperform the linear model consistently across folds and firing rate regimes. **B, C** Analysis analogous to A but for Emx1-f, Emx1-s, respectively.

**Figure S4.**
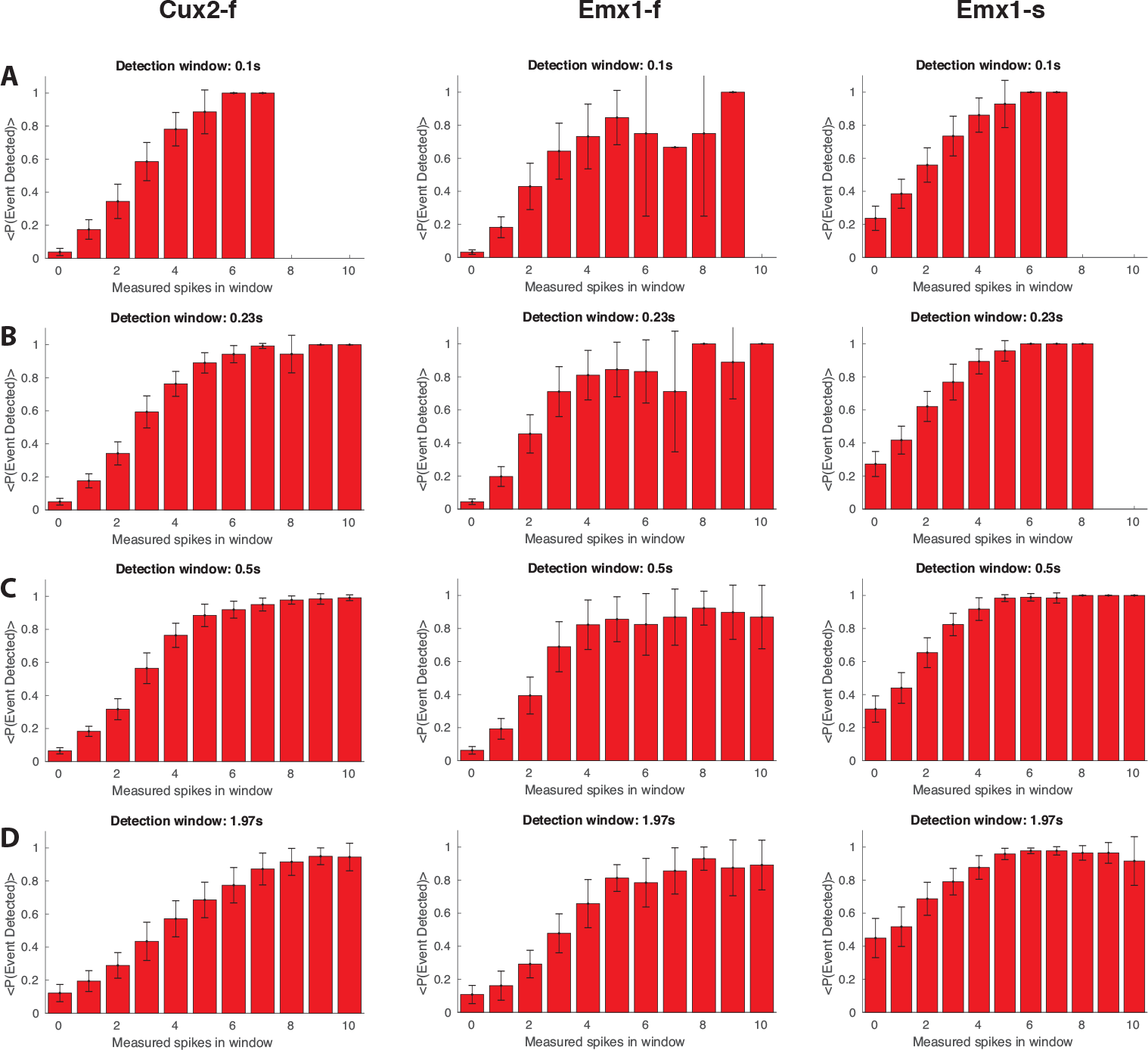
Comparison of event detection probabilities for MLSpike. In order to show that the results presented in Fig. 5 are largely independent of the employed event detection, in this figure we regenerate the results of Fig. 5 using the biophysically inspired MLSpike [19] (instead of *l*_0_-event detection). MLSpike yields fairly similar event detection probabilities as a function of the number of electrophysiologically recorded spikes for a given time window, with some minor differences in the trade-off between type 1 and type 2 errors, i.e. the single spike detection probability is higher by, approximately, the false positive detection probability.

**Figure S5.**
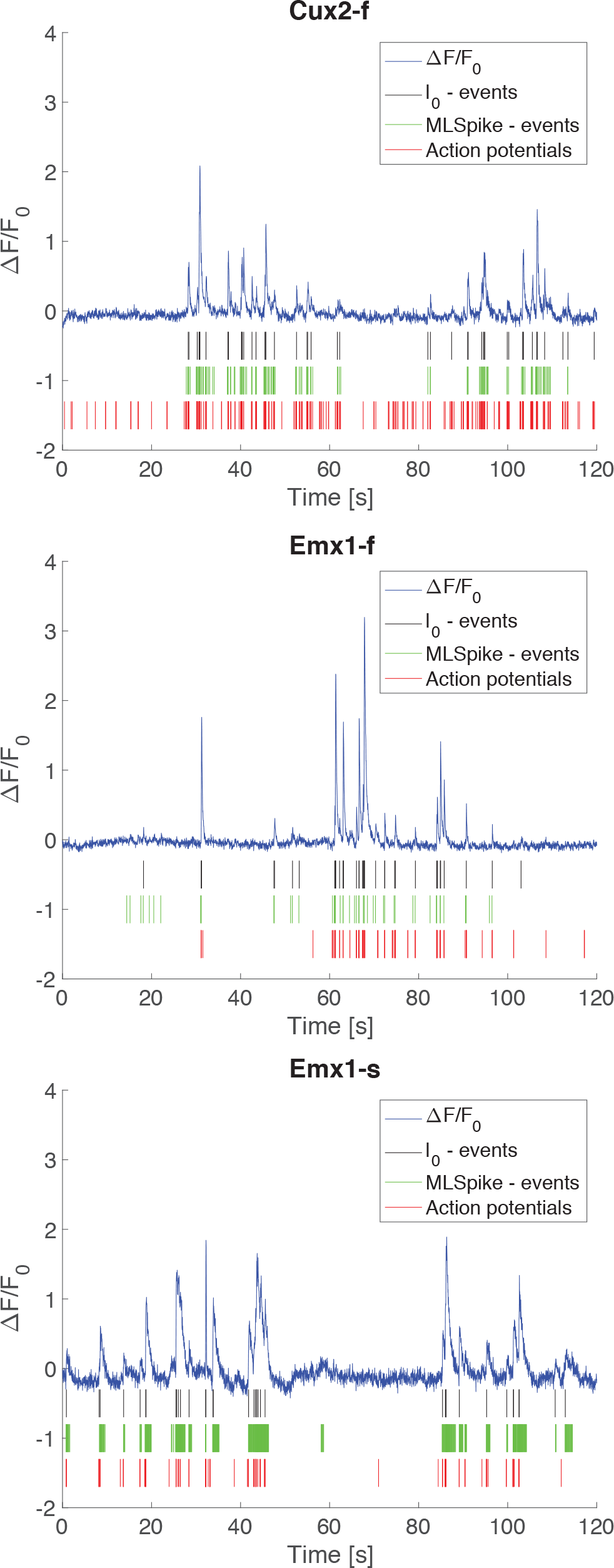
Event detection comparison. Examples of events detected by the *l*_0_ method [18] compared to events detected by MLSpike illustrate that the results are similar for these two algorithms. In the examples shown, especially for GCaMP6s, MLSpike produces more false-positives, and fewer false-negatives, consistent with Suppl. Fig. S4.

**Figure S6.**
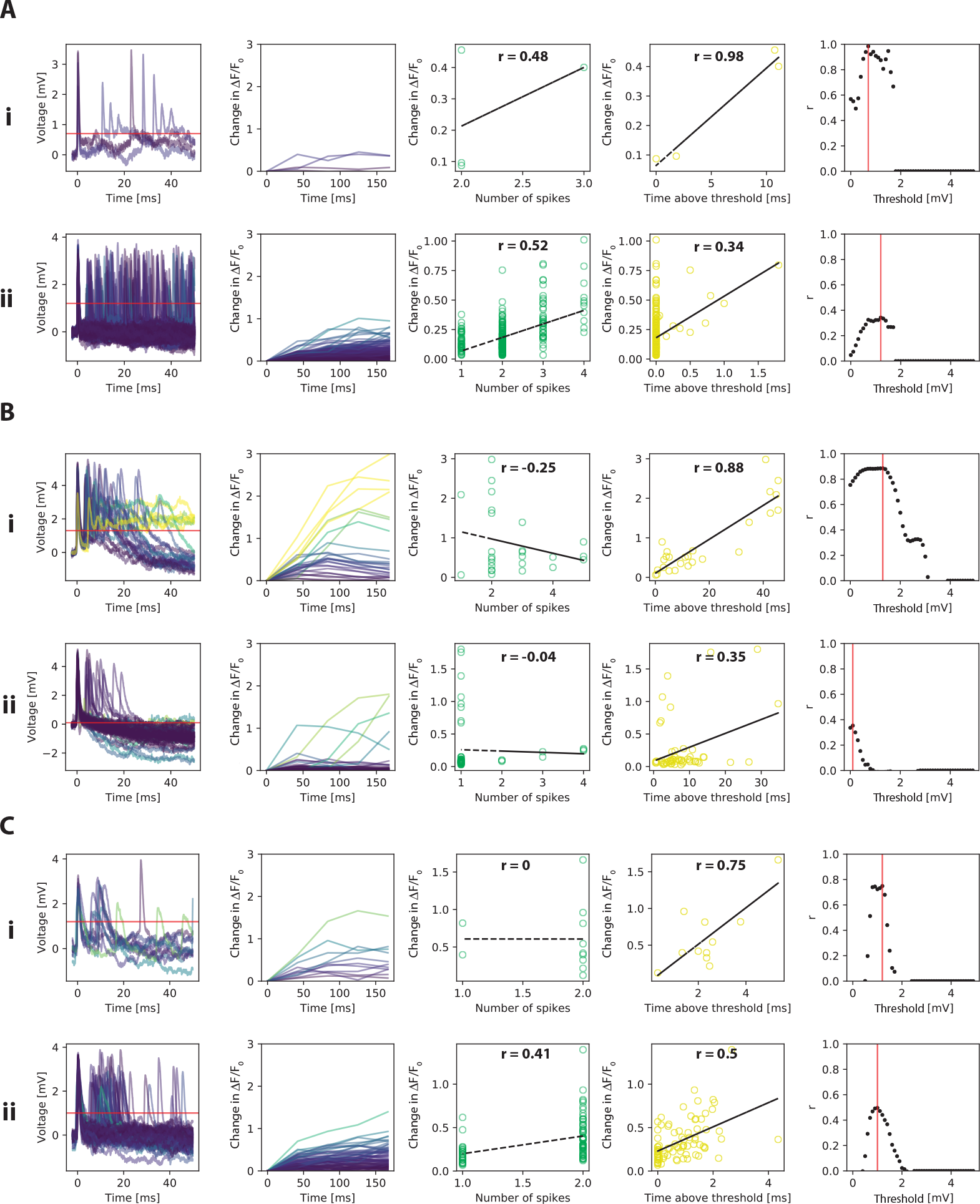
Prolonged Afterdepolarizations (PDE). **A** Data for Cux2-f i: Analysis of events with detected PDEs; from left to right: 50 ms snippets of cell-attached voltage traces; the detected spike time of the first spike is aligned at *t* = 0*s*; Change in Δ*F F*_0_ over 150 ms following the electrophysiological event (color-coded to reflect response amplitude, and to match the corresponding voltage trace); a scatter plot and linear regression (correlation coefficient *r* indicated) of the number of spikes detected in the voltage snippet against the fluorescence response (green); a scatter plot with linear regression (yellow) of the PDE duration (time the voltage trace remains above the threshold shown in red) versus the fluorescence response. For events with PDEs, time-above-threshold was generally better at explaining variance in the fluorescence response than then number of spikes preceding the event; the last scatter plot on the right shows the threshold-dependence of *r*. ii: Same analysis as in i but for events where no PDEs were detected. **B** Analysis analogous to part A but for Emx1-f, which displayed the most abundant and prominent PDEs. **C** Analysis analogous to part A but for Emx1-s.

**Figure S7.**
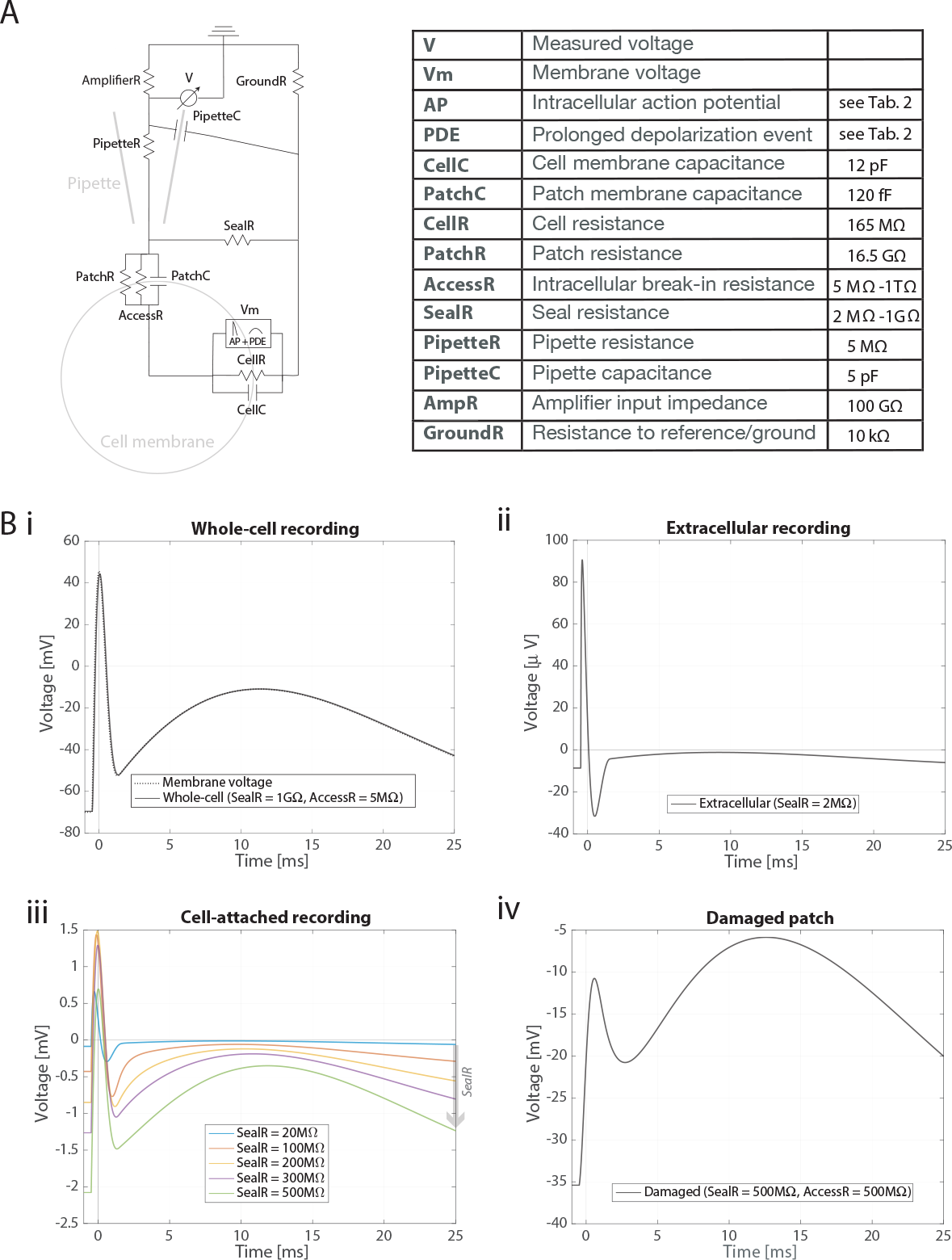
Lumped-element SPICE simulation of different patching edge cases. **A** Equivalent circuit model; the complete list of lumped element names is unpacked in the table on the right. **B** Simulation results; all waveforms were time-aligned such that the maximum membrane of the membrane potential is at *t* = 0. (i) Whole-cell recording: The membrane potential transient due to an action potential with following PDE is approximated as the sum of two chirps with biologically realistic frequency and amplitude content (black dotted line). Complete perforation of the patched membrane (AccessR = 5MΩ) results in a whole cell recording where the transient’s amplitude and time course (black line) is almost identical to that of the membrane potential. (ii) Seal resistances smaller than the pipette resistance (5 MΩ) result in transients that look like classic extracellular recordings (i.e. a derivative of the membrane voltage with amplitudes in the hundreds of *μ*V range, which peaks earlier than the membrane potential, and is ≈ 0) when the membrane potential is maximal. In the extracellular case, the APD is completely high-pass filtered out. (iii) The minimal seal resistance required for data collection (20 MΩ) yields higher-amplitude transients (on the order of 2mV peak-to-peak), which, otherwise, are reminiscent of extracellular recordings. As SealR grows larger than 100 MΩ, the transient amplitude keeps growing (¿3 mV as in most of the data presented), and the transients recorded come increasingly in-phase with the membrane potential (more reminiscent of a whole cell recording). When the Seal resistance iwas between 100 MΩ and 500 MΩ, PDEs were clearly discernible. (iv) Even small perforations of the patched membrane (1% of the parched area; AccessR = 500MΩ) result in high-amplitude transients (in excess of 20 mV), a significant lag of the transients’ time course, and a dramatic exaggeration of the PDE.

## References

1. Ian H Stevenson and Konrad P Kording. How advances in neural recording affect data analysis. Nature Neuroscience, 14(2):139–142, feb 2011.

2. J. Z. Young. STRUCTURE OF NERVE FIBRES AND SYNAPSES IN SOME INVERTEBRATES. Cold Spring Harbor Symposia on Quantitative Biology, 4(0):1–6, jan 1936.

3. Howard J. Curtis and Kenneth S. Cole. Membrane action potentials from the squid giant axon. Journal of Cellular and Comparative Physiology, 15(2):147–157, apr 1940.

4. A. L. Hodgkin and A. F. Huxley. Action Potentials Recorded from Inside a Nerve Fibre. Nature, 144(3651):710–711, oct 1939.

5. Shy Shoham, Daniel H. O’Connor, and Ronen Segev. How silent is the brain: is there a “dark matter” problem in neuroscience? Journal of Comparative Physiology A, 192(8):777–784, aug 2006.

6. Maria Göppert. Über die Wahrscheinlichkeit des Zusammenwirkens zweier Lichtquanten in einem Elementarakt. Die Naturwissenschaften, 17(48):932–932, nov 1929.

7. Maria Göppert-Mayer. Über Elementarakte mit zwei Quantensprüngen. Annalen der Physik, 401(3):273–294, jan 1931.

8. W Denk, J H Strickler, and W W Webb. Two-photon laser scanning fluorescence microscopy. Science (New York, N.Y.), 248(4951):73–6, apr 1990.

9. Tsai-Wen Chen, Trevor J. Wardill, Yi Sun, Stefan R. Pulver, Sabine L. Renninger, Amy Baohan, Eric R. Schreiter, Rex A. Kerr, Michael B. Orger, Vivek Jayaraman, Loren L. Looger, Karel Svoboda, and Douglas S. Kim. Ultrasensitive fluorescent proteins for imaging neuronal activity. Nature, 499(7458):295–300, jul 2013.

10. Saskia E J de Vries, Jerome Lecoq, Michael A Buice, Peter A Groblewski, Gabriel K Ocker, Michael Oliver, David Feng, Nicholas Cain, Peter Ledochowitsch, Daniel Millman, Kate Roll, Marina Garrett, Tom Keenan, Leonard Kuan, Stefan Mihalas, Shawn Olsen, Carol Thompson, Wayne Wakeman, Jack Waters, Derric Williams, Chris Barber, Nathan Berbesque, Brandon Blanchard, Nicholas Bowles, Shiella Caldejon, Linzy Casal, Andrew Cho, Sissy Cross, Chinh Dang, Tim Dolbeare, Melise Edwards, John Galbraith, Nathalie Gaudreault, Fiona Griffin, Perry Hargrave, Robert Howard, Lawrence Huang, Sean Jewell, Nika Keller, Ulf Knoblich, Josh Larkin, Rachael Larsen, Chris Lau, Eric Lee, Felix Lee, Arielle Leon, Lu Li, Fuhui Long, Jennifer Luviano, Kyla Mace, Thuyanh Nguyen, Jed Perkins, Miranda Robertson, Sam Seid, Eric Shea-Brown, Jianghong Shi, Nathan Sjoquist, Cliff Slaughterbeck, David Sullivan, Ryan Valenza, Casey White, Ali Williford, Daniela Witten, Jun Zhuang, Hongkui Zeng, Colin Farrell, Lydia Ng, Amy Bernard, John W Phillips, R Clay Reid, and Christof Koch. A large-scale, standardized physiological survey reveals higher order coding throughout the mouse visual cortex. Nature Neuroscience, in press 2019.

11. Bruce P. Bean. The action potential in mammalian central neurons. Nature Reviews Neuroscience, 8(6):451–465, jun 2007.

12. Lauren M. Barnett, Thomas E. Hughes, and Mikhail Drobizhev. Deciphering the molecular mechanism responsible for GCaMP6m’s Ca2+−dependent change in fluorescence. PLOS ONE, 12(2):e0170934, feb 2017.

13. Lawrence Huang, Ulf Knoblich, Peter Ledochowitsch, Jack Waters, Jerome Lecoq, Clay R. Reid, Christof Koch, Saskia de Vries, Michael A. Buice, Gabe Murphy, Hongkui Zeng, and Lu Li. Relationship between in vivo calcium events and spiking activity in transgenic mouse lines. bioRxiv, 2019.

14. Siegfried Weisenburger, Frank Tejera, Jeffrey Demas, Brandon Chen, Jason Manley, Fraser T Sparks, Francisca Martínez Traub, Tanya Daigle, Hongkui Zeng, Attila Losonczy, and Alipasha Vaziri. Volumetric Ca2+ Imaging in the Mouse Brain Using Hybrid Multiplexed Sculpted Light Microscopy. Cell, 177(4):1050–1066.e14, may 2019.

15. R Quian Quiroga, Z Nadasdy, and Y Ben-Shaul. Unsupervised spike detection and sorting with wavelets and superparamagnetic clustering. Neural computation, 16:1661–1687, 2004.

16. Martin Ester, Martin Ester, Hans-Peter Kriegel, Jörg Sander, and Xiaowei Xu. A density-based algorithm for discovering clusters in large spatial databases with noise. pages 226—-231, 1996.

17. Erich Schubert, Jörg Sander, Martin Ester, Hans Peter Kriegel, and Xiaowei Xu. DBSCAN Revisited, Revisited. ACM Transactions on Database Systems, 42(3):1–21, jul 2017.

18. Sean Jewell, Toby Dylan Hocking, Paul Fearnhead, and Daniela Witten. Fast Nonconvex Deconvolution of Calcium Imaging Data. feb 2018.

19. Thomas Deneux, Attila Kaszas, Gergely Szalay, Gergely Katona, Tamás Lakner, Amiram Grinvald, Balázs Róozsa, and Ivo Vanzetta. Accurate spike estimation from noisy calcium signals for ultrafast three-dimensional imaging of large neuronal populations in vivo. Nature Communications, 7:12190, jul 2016.

20. Joshua T. Vogelstein, Adam M. Packer, Timothy A. Machado, Tanya Sippy, Baktash Babadi, Rafael Yuste, and Liam Paninski. Fast Nonnegative Deconvolution for Spike Train Inference From Population Calcium Imaging. Journal of Neurophysiology, 104(6):3691–3704, dec 2010.

21. Johannes Friedrich, Pengcheng Zhou, and Liam Paninski. Fast online deconvolution of calcium imaging data. PLOS Computational Biology, 13(3):e1005423, mar 2017.

22. Artur Speiser, Jinyao Yan, Evan W Archer, Lars Buesing, Srinivas C Turaga, and Jakob H Macke. Fast amortized inference of neural activity from calcium imaging data with variational autoencoders. In I Guyon, U V Luxburg, S Bengio, H Wallach, R Fergus, S Vishwanathan, and R Garnett, editors, Advances in Neural Information Processing Systems 30, pages 4024–4034. Curran Associates, Inc., 2017.

23. Marius Pachitariu, Carsen Stringer, and Kenneth D Harris. Robustness of Spike Deconvolution for Neuronal Calcium Imaging. The Journal of neuroscience: the official journal of the Society for Neuroscience, 38(37):7976–7985, sep 2018.

24. David S Greenberg, Damian J Wallace, Kay-Michael Voit, Silvia Wuertenberger, Uwe Czubayko, Arne Monsees, Takashi Handa, Joshua T Vogelstein, Reinhard Seifert, Yvonne Groemping, and Jason ND Kerr. Accurate action potential inference from a calcium sensor protein through biophysical modeling. bioRxiv, page 479055, nov 2018.

25. Nicholas A Steinmetz, Christina Buetfering, Jerome Lecoq, Christian R Lee, Andrew J Peters, Elina A K Jacobs, Philip Coen, Douglas R Ollerenshaw, Matthew T Valley, Saskia E J de Vries, Marina Garrett, Jun Zhuang, Peter A Groblewski, Sahar Manavi, Jesse Miles, Casey White, Eric Lee, Fiona Griffin, Joshua D Larkin, Kate Roll, Sissy Cross, Thuyanh V Nguyen, Rachael Larsen, Julie Pendergraft, Tanya Daigle, Bosiljka Tasic, Carol L Thompson, Jack Waters, Shawn Olsen, David J Margolis, Hongkui Zeng, Michael Hausser, Matteo Carandini, and Kenneth D Harris. Aberrant Cortical Activity in Multiple GCaMP6-Expressing Transgenic Mouse Lines. eNeuro, 4(5), 2017.

